# Episomal editing of synthetic constructs in yeast using CRISPR

**DOI:** 10.1101/2022.06.21.496881

**Authors:** Yu Zhao, Camila Coelho, Stephanie Lauer, Jon M. Laurent, Ran Brosh, Jef D. Boeke

## Abstract

Use of synthetic genomics to design and build “big” DNA has revolutionized our ability to answer fundamental biological questions by employing a bottom-up approach. *S. cerevisiae*, or budding yeast, has become the major platform to assemble large synthetic constructs thanks to its powerful homologous recombination machinery and the availability of well-established molecular biology techniques. However, efficiently and precisely introducing designer variations to episomal assemblies remains challenging. Here, we describe CRISPR Engineering of EPisomes in Yeast, or CREEPY, for rapid engineering of mammalian DNA constructs larger than 100 kb. We demonstrate that editing of circular episomes presents unique challenges compared to modifying native yeast chromosomes with CRISPR. After optimizing CREEPY for episomal editing, we achieve efficient simplex and multiplex editing as demonstrated by engineering a mouse *Sox2*-harboring episome.

## Introduction

Synthetic biology approaches have been used to redesign and assemble “big” DNA to generate chromosomes and entire genomes, including *de novo* assembly of synthetic poliovirus and bacteriophage DNA, as well as the prokaryotic genomes of *Mycoplasma* and *E. coli*^1, 2, 3, 4^. In eukaryotes, the Sc2.0 project aims to build synthetic chromosomes in yeast from the bottom up^5, 6, 7, 8^. As technology rapidly advances, genome writing in mammalian cells has also become feasible, resulting in new large DNA manipulation and delivery strategies at the locus level^9, 10, 11^. These studies provide a new lens to understand complex genome architecture, expression regulation, and the genetic basis of human disease.

The human genome, like other mammalian genomes, is complex, with introns and non-coding sequences such as repeats and regulatory elements. Furthermore, most human genome variants implicated in disease map to non-coding, regulatory regions^12^. Recently, synthetic regulatory reconstitution has been used to map the regulatory architecture of the *HoxA* cluster by ectopically integrating assembled rat *HoxA* cluster variants in mouse embryonic stem cells (mESCs)^13^. Engineering such variations as part of big DNA assembly is critical for understanding their role and regulation within their native context.

The assembly of large (>100 kb) DNA constructs in *S. cerevisiae* is a fundamental step towards writing mammalian genomes. Yeast is tractable as a platform to assemble large DNA constructs because of its efficient homologous recombination (HR) machinery and the advanced molecular toolset available to researchers. Using HR of overlapping segments, linear DNA fragments can be assembled as large circular episomal yeast and bacterial artificial chromosomes (YAC/BAC) constructs that can be propagated and transferred to bacteria for isolation before delivering to mammalian cells. After assembly, constructs can be modified in yeast to generate panels of designer variants. Compared to building many different constructs from scratch, it is also much more feasible and reliable to introduce designer modifications into a parental base assembly. Episomal constructs can be delivered to mammalian systems, or further characterized in yeast, as *S. cerevisiae* is widely used to optimize episomal biosynthetic pathways in metabolic engineering of natural products^14, 15, 16, 17^.

CRISPR has been widely used for yeast genome editing^18, 19^. Directed by a sequence-specific single guide RNA (sgRNA), the Cas9 nuclease creates a DNA double strand break (DSB). This DSB can be repaired by homologous recombination with a co-transformed donor DNA containing polymorphisms that prevent further Cas9 binding and cleavage to achieve successful editing. In the absence of such a donor DNA template, the original genomic break may lead to cell cycle arrest or the loss of an essential gene and subsequent death. Compared to mammalian cells, *S. cerevisiae* performs non-homologous end joining (NHEJ) with high fidelity, with the major repair product being simple religation^20^. As a result, the repaired DNA reforms the original Cas9 cleavage site unless errors in ligation such as lost bases prevent further Cas9 recognition. Typically, when CRISPR/Cas9 targets a chromosomal site that is efficiently cut, the number of surviving colonies in the absence of donor template DNA are 100-to 1000-fold lower than in its presence^18^. Previous CRISPR studies have focused mainly on genomic editing, with many well-designed systems established for both simplex and multiplex targets^21, 22, 23^. However, the efficiency of editing episomal DNA constructs by CRISPR/Cas9 remains unclear and a CRISPR toolbox especially optimized for episomes is lacking. It is also unknown whether there are any fundamental differences between episomal and chromosomal editing in yeast.

In this study, we introduce CRISPR Engineering of EPisomes in Yeast, or CREEPY, for episomal DNA engineering. We first compare the efficiency of CRISPR/Cas9 for targeting episomes and chromosomes. With CREEPY optimized for episomal editing, we achieve simplex and multiplex editing, as demonstrated by engineering of a 143-kb *mSox2* assembly containing the mouse *Sox2* gene and regulatory regions^24^. *Sox2* is a Yamanaka factor essential for maintaining stem cell pluripotency^25^. The *mSox2* episome represents a typical big DNA assembly in length (>100 kb) and contains the same complex mammalian genomic elements as the native locus. While the majority of episomal edits are successful, unintended modifications can occur. We identified the mechanism underlying these modifications, which occur when internal deletions result from end joining of microhomologous repeats flanking the initial DSB site rather than errors from NHEJ. The CREEPY constructs and methods developed here can be used as a resource to further advance the fields of DNA assembly and metabolic engineering in yeast and mammalian systems.

## Results

### CRISPR/Cas9 constructs and genomic editing in yeast

To determine whether there are any distinct challenges for episomal editing compared to standard chromosomal editing, we developed a CRISPR system based on previously tested constructs^18^. In this study, we used a human codon-optimized *S. pyogenes* Cas9 driven by the yeast *TEF1* promoter together with a *CYC1* terminator in a *CEN/ARS* vector (pCTC019) (Fig. 1a). In a two-plasmid system, the sgRNA is expressed using the RNA polymerase III (pol III) *SNR52* promoter and terminated by the poly T sequence in the *SUP4* terminator on a separate plasmid (pNA0306). The parent strain is pre-transformed with the Cas9 plasmid, followed by a second transformation of the sgRNA plasmid with donor DNA. We also subcloned both Cas9 and sgRNA expression modules into one single plasmid, with either a *CEN/ARS* (pYZ462) or *2μ* (pYZ463) backbone, using the same promoters and terminators for Cas9 (Fig. 1b). This plasmid enables delivery of all the components required for editing in a single transformation step, thereby increasing the throughput of CRISPR editing.

**Fig. 1.**
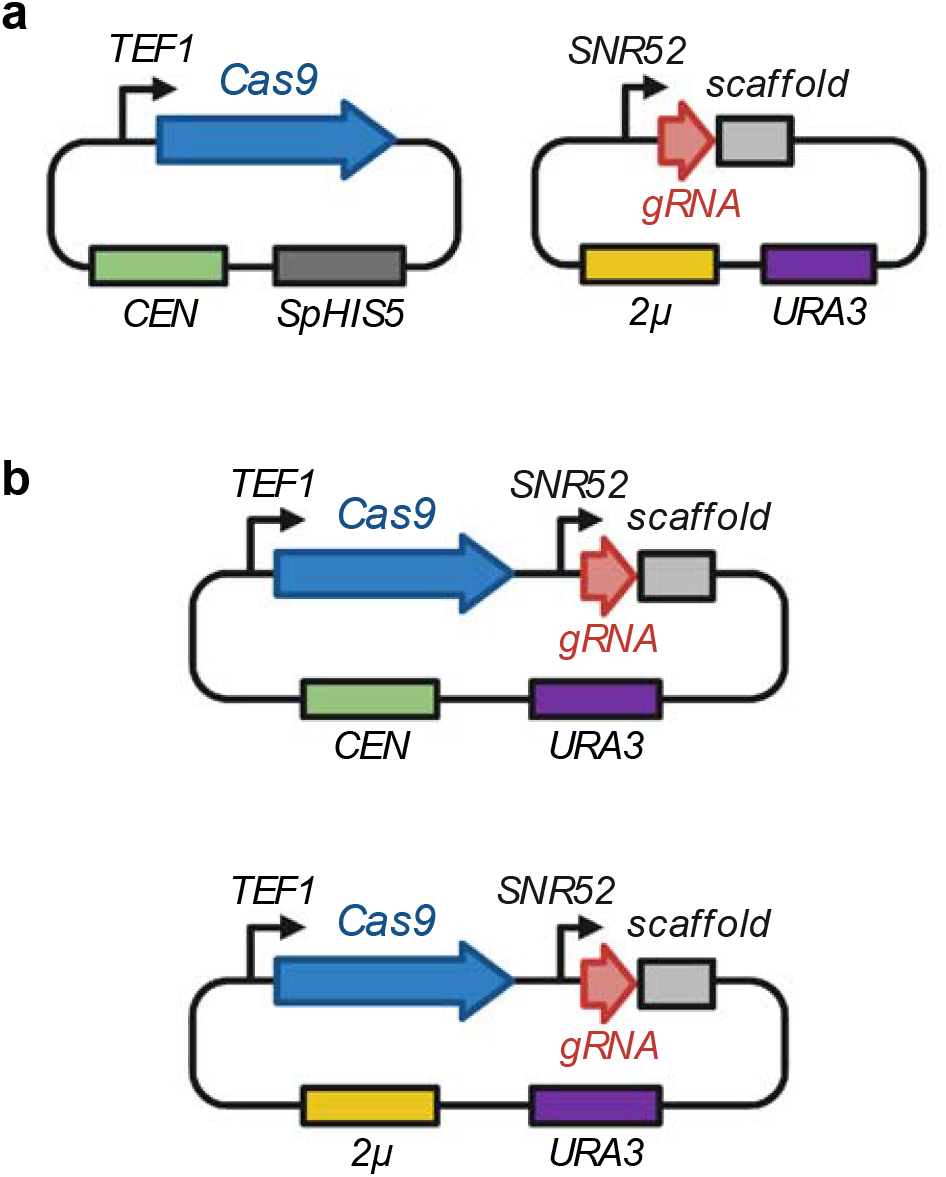
CRISPR/Cas9 constructs used in this study. **a**. The two-plasmid system. Cas9 plasmid is pre-transformed into yeast, followed by a second transformation of the sgRNA plasmid. **b**. The single-plasmid system. Cas9 and sgRNA are expressed from a single plasmid, with *CEN/ARS* (up) or *2μ* backbone (bottom).

To compare episomal editing to genome editing, we first targeted the *ADE2* coding sequence (CDS) on yeast chromosome *XV* using a previously reported sgRNA^23^. This sgRNA was assembled into three constructs (two-plasmid, and single-plasmid with *CEN/ARS* or *2μ* backbone). The constructs were delivered to BY4741 together with an appropriate donor template DNA and red colonies on transformation plates, indicating successful deletion of *ADE2*, were counted to estimate editing efficiencies. All three systems achieved efficient editing, with a >97% success rate (Fig. 2a). Notably, the two-plasmid system in which the Cas9 plasmid is transformed prior to introducing the guide RNA showed the highest efficiency, though the difference was not significant, suggesting that pre-transformed abundant Cas9 in cells may maximize genomic editing efficiency. Despite the slightly reduced efficiency of the single-plasmid systems, the convenience offered by a single transformation makes them a more attractive option. Transforming the Cas9/sgRNA without a donor DNA template resulted in many fewer colonies (∼1%) (Fig. 2b), confirming previous reports that yeast cells largely rely on HR with donor templates or high-fidelity NHEJ to repair DSBs.

**Fig. 2.**
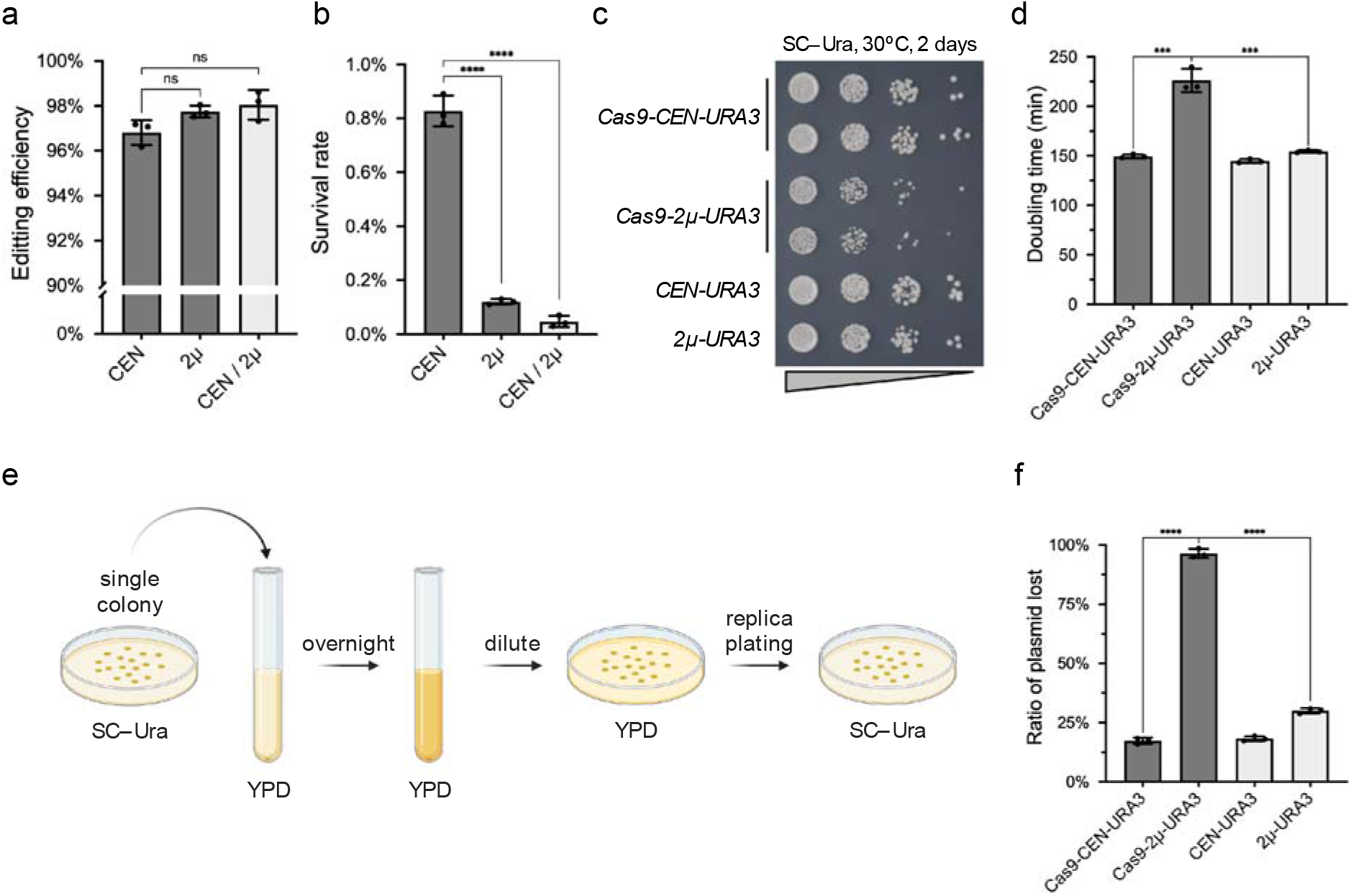
Genomic editing in yeast with one- or two-plasmid systems. **a**. Editing efficiency targeting genomic *ADE2*, as determined by three replicates. Error bars represent mean ± SD. ns, not significant as analyzed with unpaired t test. **b**. Survival rate of colonies transformed with Cas9/sgRNA alone without donor DNA. Error bars represent mean ± SD, n=3 and p<0.0001 (****), as analyzed with unpaired t test. **c**. Spot assay on SC–Ura plates with BY4741 strains carrying Cas9 expressed from *CEN/ARS* or *2μ* plasmids. The pRS416 (*CEN-URA3*) and pRS426 (*2μ-URA3*) are used as empty plasmid controls. **d**. Doubling time of the same yeast strains used in the spot assay. Error bars represent mean ± SD, n=3 and p<0.001 (***), as analyzed with unpaired. Growth curves are shown in Supplementary Figure 1. **e**. Method to test the Cas9 toxicity and plasmid stability. **f**. Proportion of colonies lost with *URA3* plasmid. Error bars represent mean ± SD, n=3 and p<0001 (****), as analyzed with unpaired t test.

While abundant Cas9 protein presumably increases chromosomal editing efficiency, we considered that the Cas9 load may also lead to toxicity in yeast. To assess this directly, we transformed the same Cas9 expression module with either the *CEN/ARS* or *2μ* origins, respectively, and then performed a spot assay to determine strain fitness (Fig. 2c). Compared to wild-type cells with empty pRS416 and pRS426 vectors, the Cas9-CEN construct had a minor effect on cell growth, while the Cas9-2μ construct showed obvious toxicity. Growth of these strains in liquid selective medium showed results consistent with our spot assay (Supplementary Figure 1). The doubling time of the strain with the Cas9-2μ plasmid was significantly longer (∼1.5×) compared to the strains with Cas9-CEN or empty vectors (Fig. 2d). These results indicate that Cas9 expressed from the high-copy *2μ* plasmid leads to obvious toxicity associated with increased Cas9 abundance.

In practice, CRISPR plasmids should be removed after editing as soon as possible in order to minimize off-target effects. Plasmid loss also enables repeated use of auxotrophic marker genes for further editing or experimenting. Cas9 toxicity and plasmid stability may affect this process. To test this, we inoculated single colonies transformed with either the Cas9-CEN or Cas9-2μ plasmid into non-selective liquid media and then estimated the ratio of cells that lost the plasmid by replica plating (Fig. 2e). To provide a baseline measurement, we also determined plasmid loss rates of an empty *2μ* plasmid (pRS426) and an empty *CEN/ARS* plasmid (pRS416). As expected, the empty *2μ* plasmid was lost at a higher rate than the empty *CEN/ARS* plasmid (Fig. 2f)^26^. Consistent with the toxicity of the Cas9 protein reported above, the high-copy Cas9-2μ plasmid was lost at an even higher rate than the empty *2μ* plasmid. The Cas9-CEN construct showed a similar loss ratio compared to the empty CEN/ARS plasmid, consistent with minimal toxicity to yeast cells.

### CRISPR/Cas9 engineering of episomes in yeast - targeting *mSox2* assembly

To test the editing efficiency with CREEPY, we used an episomal construct containing 143-kb wild-type mouse *Sox2* (*mSox2*) fragment, which includes the coding sequence and distal regulatory clusters such as DNase I hypersensitive sites (DHSs) and CTCF-binding sites (Fig. 3a)^24^. First, we tested episomal editing by deleting a single CTCF site, CTCF8, in the *mSox2* construct (Fig. 3b). The sgRNA was assembled into the same constructs as described above, and the same number of plasmids and donor templates were used as in chromosomal editing. Following selection of transformants, single colonies were screened using colony PCR for successful designer deletion of CTCF8 using deletion-specific primers (Supplementary Figure 2). We found the Cas9/sgRNA-CEN construct showed the lowest efficiency (∼70%), while the Cas9/sgRNA-2μ construct and two-plasmid system had high editing efficiency (∼96%) (Fig. 3c). This suggests that high Cas9 protein abundance, either from pre-transformed Cas9 plasmid or the high-copy *2μ* construct, is important for efficient episomal editing.

**Fig. 3.**
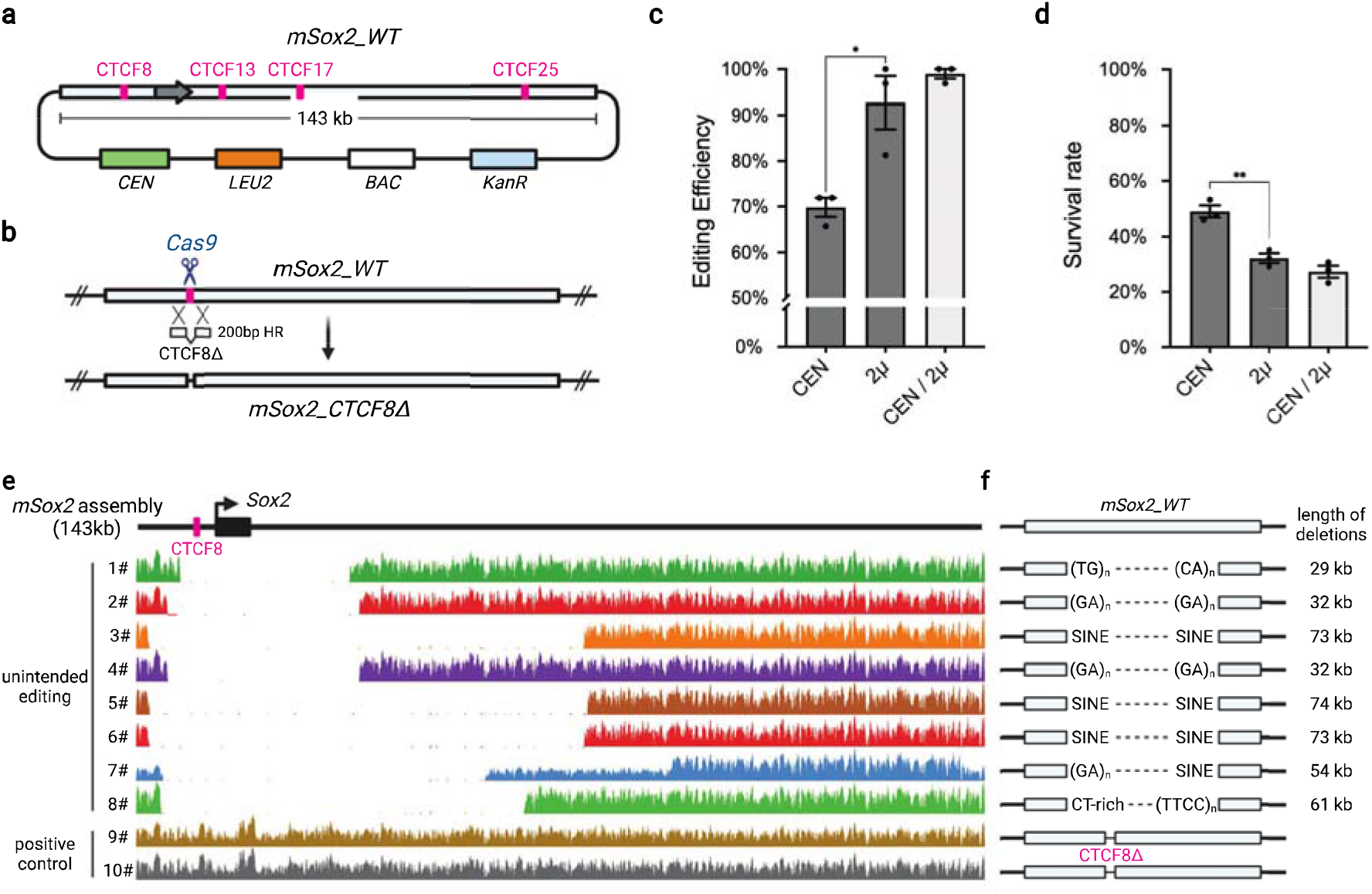
Episomal editing in yeast targeting at mSox2 assembly. **a**. Structure of episomal mSox2 assembly. **b**. Episomal editing strategy. Cas9 creates a DSB at CTCF8 which can be repaired with a donor DNA containing CTCF8Δ. **c**. Editing efficiency with one-plasmid (CEN or 2μ backbone) or two-plasmid system (CEN/2μ) as determined by genotyping using deletion-specific primers (Supplementary Figure 2). Error bars represent mean ± SD. n=3 and p<0.05 (*), as analyzed with unpaired t test. **d**. Survival rate of colonies transformed with Cas9/gRNA target CTCF8 without donor DNA templates. Error bars represent mean ± SD. n=3 and p<0.005 (**), as analyzed with unpaired t test. WGS read coverage of colonies with unintended or successful edits aligned to the mSox2 assembly. **f**. Unintended deletion boundaries from mapped to the indicated mouse genome repetitive elements (see also Supplementary Table 1).

Considering all these results, we determined that best system for **genomic** editing where the priority is usually to build a yeast strain with designer modifications quickly and efficiently, and avoid non-specific mutations, is the single-plasmid system containing Cas9 on the *CEN/ARS* backbone, especially given its minor toxicity, minimal effect on cell growth, and efficient loss after culturing in non-selective medium (Fig. 2f). In contrast, **episomal editing** is primarily used for generating DNA variants and thus Cas9 on the *2μ* backbone usually provides the best option because of its higher efficiency and rapid loss rate. In the following experiments, the Cas9-2μ construct was used.

### Unintended episomal edits are due to MMEJ between microhomologous repeats

For chromosomal editing, very few colonies (∼1%) appeared in the absence of donor DNA (Fig. 2b). In contrast, episomal targeting by Cas9 in the absence of donor DNA resulted in a much higher survival rate (∼40%, Fig. 3d), suggesting that a different mechanism might be used to recover from DSBs in episomes than in chromosomes. An alternative explanation is that the repair mechanisms are similar, but the selective pressures involving episomes and chromosomes, which contain many essential genes, are very different. Moreover, the mammalian DNA is richer in repeats compared with yeast genomic DNA. To uncover the mechanism responsible for the background in the absence of donor DNA, we selected 8 independent colonies from the transformation that did not pass PCR screening for whole genome sequencing (WGS). Two colonies with successful CTCF8 deletion were used as positive controls. WGS reads were aligned to the *mSox2* 143-kb construct and the Cas9/gRNA-2μ plasmids references (Fig. 3e and Supplementary Figure 3-4). First, CTCF8 deletion was confirmed in the positive controls, which were otherwise intact (Supplementary Figure 3). However, we found large internal deletions (30-70kb) that always included the original DSB site in the other eight samples that failed genotyping. These deletions allowed cells to avoid repeated digestion of the episomes by Cas9. This result is dramatically different compared to chromosomal editing, in which survivors tend to present point mutants at the site of cleavage^18^. The Cas9/gRNA construct remained intact in all of these samples (Supplementary Figure 4), indicating that the CRISPR system was still functionally active.

**Fig. 4.**
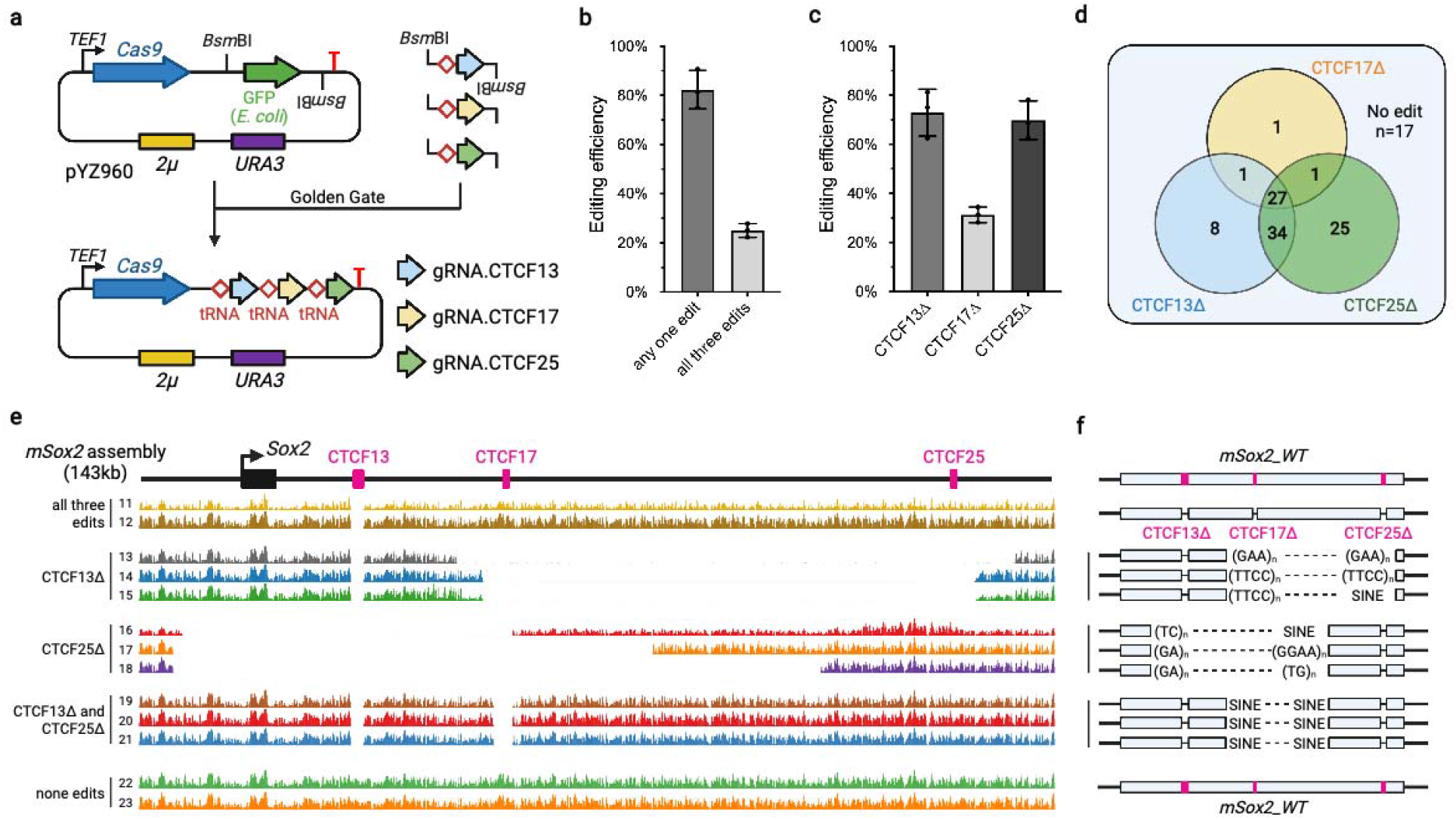
Multiplex editing for more CTCF sites in mSox2 assembly. **a**. One-plasmid system for multiple episomal editing. The tRNA-gRNA array is built using Golden Gate assembly with BsmBI. **b**. Episomal editing efficiency for at least one or all three designer deletions. Error bars represent mean ± SD (n=3). **c**. Episomal editing efficiency calculated for each designer deletion. Error bars represent mean ± SD (n=3). **d**. Distributions of successful edits of CTCF13Δ, CTCF17Δ and/or CTCF25Δ. **e**. WGS read coverage from colonies obtained in multiplex episomal editing experiments aligned to the mSox2 assembly. **f**. Genomic boundaries of unintended deletions from multiple episomal editing experiments mapped to the mouse genome and annotated with relevant repetitive elements mapping at the sites of deletion (see also Supplementary Table 2).

Next, we aligned the WGS reads to the mm10 genome and inspected the sequence properties adjacent to the deletion boundaries using RepeatMasker^27^. Although the boundaries of deletions were variable, we found they all mapped to repetitive elements, including microsatellite, SINE and low complexity repeats (Fig. 3f). These repetitive sequences are present in the wild-type mouse genome and as such were an appropriate part of the original *mSox2* assembly (Supplementary Table 1). Considering their length and composition, the deletions were most likely formed by microhomology-mediated end joining (MMEJ), rather than NHEJ or HR. In yeast cells, MMEJ is initiated by DNA resections from the DSB end, and subsequent microhomologous annealing^28^. It is an error-prone repair mechanism that is associated with deletions flanking the DSB site^20^. In contrast, the products of *S. cerevisiae* NHEJ are generally precise religations back to the wild-type sequence and HR is generally error free^28^. The observed deletions were also much longer than regular indels from non-specific NHEJ. Thus, we speculated that MMEJ occurred after Cas9 cleavage and led to the survival of cells carrying episomal constructs with internal deletions.

When transformed with Cas9/gRNA-2μ plasmids and donor DNA, yeast cells demonstrated a high efficiency of simplex episome editing efficiency (∼90%) with little background of unintended editing (∼10%). In the absence of donor DNA, survival rate is relatively high (∼40%), suggesting that yeast cells use HR to repair DSBs when donor DNA is available and that MMEJ is a secondary response when HR is not possible.

### Multiplex episomal editing using a tRNA-gRNA array

Multiplex editing will significantly accelerate introduction of more than one designer variation in parallel. With this goal in mind, we evaluated the efficiency of multiplex editing of episomes. With abundant Cas9 enzyme in cells, a key challenge is whether more than one sgRNA can be expressed at once. In yeast, several genomic editing strategies have been reported where sgRNA maturation is assisted by virus ribozymes, Csy4 cleavage or tRNA processing^19, 22, 23^. In this study, we used a tRNA-gRNA array to express multiple sgRNAs simultaneously (Fig. 4a). The sgRNAs are co-transcribed with a tRNA by pol III, and are then released from the primary transcript by endogenous RNase P and RNase Z^23, 29^. This design ensures the co-expression of multiple gRNAs within one single cell and avoids complicated strain engineering such as integration of Csy4^22^. Recently, it was also confirmed by RNA-seq that this strategy results in even expression of each gRNA from any position in the same type of array^30^.

To test multiplex editing of *mSox2*, we attempted to delete three different CTCF sites in parallel: CTCF13, CTCF17 and CTCF25 (Fig. 3a). We first built an entry vector with the same Cas9 expression module and *2μ* plasmid backbone, and containing *Bsm*BI sites for integration of the sgRNAs. Three *mSox2*-targeting sgRNAs were assembled with Golden Gate cloning (see methods) and the resulting Cas9/sgRNA-array *URA3* construct was transformed with all three corresponding donor DNAs mediating CTCF site deletions, to SC– Ura plates. First, we found that most colonies had at least one edit (∼80%), consistent with the high editing efficiency of CREEPY (Fig. 4b). More importantly, ∼25% of colonies presented all three edits, demonstrating that multiplex episomal editing was successful. We found that deletion of CTCF13 and CTCF25 showed similarly high efficiencies whereas the CTCF17 deletion was less efficient, and was mostly responsible for reducing overall multiplex editing efficiency (Fig. 4c).

To understand why some colonies had one or two but not three modifications, we investigated the distribution of these edits as shown in a Venn diagram (Fig. 4d). Most of the colonies with a successful CTCF17 deletion also contained the other two deletions (27 out of 30). In other words, as long as the CTCF17 designer deletion occurred, all three edits were likely to also have occurred. In comparison, for episomes with CTCF13Δ, the majority also contained an additional edit of CTCF25Δ (61 out of 70), but all three edits occurred in only 27 of 70 colonies. A similar ratio was observed for CTCF25Δ. Consistent with its low editing efficiency, CTCF17Δ seemed to represent the bottleneck limiting multiplex editing.

To understand the reasons behind this phenomenon, we performed WGS and aligned reads to the episomal *mSox2* and CRISPR plasmid from three independent colonies of each of the following edit types: all three edits, CTCF13Δ only, CTCF25Δ only, CTCF13Δ and CTCF25Δ, and no detected edits, as determined by PCR genotyping (Fig. 4e). For colonies with all three edits, the designer deletions were confirmed and the *mSox2* constructs were otherwise intact (Supplementary Figure 5). Interestingly, for colonies with CTCF13Δ only, in addition to validating the deletion of CTCF13, internal deletions in the downstream region flanking CTCF17 and CTCF25 were observed, completely eliminating those Cas9/sgRNA recognition sites. Similar events were observed in colonies with CTCF25Δ only, where upstream internal deletions caused the loss of CTCF13 and CTCF17 sites. Consistently, for colonies with both CTCF13Δ and CTCF25Δ but not CTCF17Δ, unintended deletions were only detected for the CTCF17 site. Notably, the CRISPR plasmids from all these groups remained intact with full-length tRNA-sgRNA arrays (Supplementary Figure 6). These results indicate that all three sgRNAs, including gRNA.CTCF17, were functional and expressed well, as cleavage happened to all CTCF sites. CTCF17 is located centrally in the *mSox2* construct while CTCF13 and CTCF25 sites are terminal. Thus, there is a larger number of repetitive elements situated on both sides of CTCF17. Also, it is notable that the CTCF17 site is flanked on both sides by mouse SINE B1 and B2 retrotransposons, most of which range in length from 150-180 bp. Due to their length, these repeats provide rather extensive opportunities for both MMEJ-mediated and HR-mediated repair and this may also contribute significantly to the relatively-high frequency of recovering this class of deletions. Rather than sgRNA performance, this “position effect” is probably the major cause of lower efficiency of CTCF17 deletion. Finally, for colonies with 0 edits, wild-type *mSox2* was observed without any modifications. Instead, in those cases, deletions occurred in the Cas9/sgRNA-array plasmid, disrupting the Cas9 CDS (Supplementary Figure 6).

We also aligned the sequencing reads to the mouse genome (mm10). Consistent with the simplex editing experiment, we mapped all the deletion boundaries to microhomologous repeats, such as micro-satellites and SINEs (Fig. 4f and Supplementary Table 2). As we speculated previously, this finding further demonstrated that yeast cells can resolve Cas9 cleavage in episomes through formation of extensive internal deletions promoted by MMEJ that destroy the cleavage sites.

## Discussion

In this study, we used CREEPY to perform CRISPR editing of episomes. CRISPR has been widely used for strain construction and genome engineering. Both the Cas9 and Cas12a systems have been implemented for genomic editing in yeast for simplex and multiplex targets, and optimized for a variety of functions^31, 32, 33, 34, 35^. Here we used Cas9 with human-optimized codons, driven by the constitutively active *TEF1* promoter, as described in its first implementation in *S. cerevisiae*^18^. Other groups have tested the same Cas9 enzyme with codons optimized for yeast or using the original coding sequence from *S. pyogenes*^19, 21, 36^. All versions of the Cas9 enzyme showed similarly high genomic editing efficiency, indicating that the effects of codon-optimization are minimal. However, more direct comparisons of efficiency and toxicity are lacking and represent one area of further study.

Here, we demonstrated that a single plasmid system with Cas9 in a *2μ* backbone led to obvious toxicity and reduced growth rate in yeast cells. This construct should be avoided for genomic editing. In other experiments unrelated to this study, we found that genome editing in haploid strains with the Cas9-2μ construct might cause whole genome endoreduplication, generating a diploid cell with successful editing but two copies of each chromosome (data not shown). Thus, we recommend using the Cas9-CEN construct to reduce off-target effects while maintaining optimal host strain fitness, i.e., for yeast genome engineering. However, for Big DNA applications focused on mammalian genome rewriting, yeast is used as a platform for DNA assembly only. The final constructs are extracted and the accompanying yeast genome is no longer of any consequence. Thus, we recommend using the Cas9-2μ construct for CREEPY due to its higher editing efficiency in this context.

To test episomal editing, we used CREEPY to delete CTCF8, CTCF13, CTCF17 and CTCF25 in the *mSox2* assembly, generating a construct with four CTCF sites removed in two steps. Beyond this, with CREEPY, over 60 constructs with deletions, inversions and surgical alterations of DHSs and CTCF of *mSox2*, were built in yeast and delivered to mESCs in order to study its regulatory architecture^24^. In practice, we also tried another single-plasmid system, pYTK-Cas9, built with the yeast tool kit (Supplementary Figure 7a)^37^. It contained a yeast-codon optimized Cas9, driven by the constitutive yeast *PGK1* promoter in a *CEN/ARS* backbone. Different auxotrophic markers for the Cas9 plasmid backbone were also used, including *URA3* (pNA0304, pYZ462 and pYZ463), *ScHIS3* (pNA0519) and *SpHIS5* (pCTC019) (Supplementary Figure 7b). All of these constructs worked efficiently. For sgRNA expression, the *SNR52* promoter and endogenous tRNA were used in this study. Other pol III promoters like *RPR1* can also be used to express sgRNAs in yeast^38, 39^. All these promoters are pol III promoters that worked well with high efficiency. The success of these experiments highlights the utility of using a Big DNA approach to study synthetic regulatory genomics. Compared to yeast episomes, the mammalian extrachromosomal circular DNA (eccDNA) is shown to be prevalent in human disease, especially in cancer ^40^. But its biological functions and regulations still need further study. We believe developing similar engineering methods or models to investigate eccDNA’s regulation in mammalian system will be of much interest.

We achieved both simplex and multiplex episomal editing in our study. Facilitated by a tRNA-gRNA array, the simultaneous expression of multiple gRNAs is very feasible. We speculate that the limiting step for episomal editing is the available copy number of each of the multiple donor DNA fragments required. During genome editing, yeast growth is arrested due to DSBs in the absence of any single donor DNA. For episomes, successful multiplex editing requires the presence of all donor DNAs in one cell. Missing one or some of these templates may lead to survival but inefficient editing as demonstrated by the multiplex editing we achieved here. This process may become increasingly challenging with additional targets and represents an avenue for further improvements to multiplex editing in yeast.

As Big DNA design and writing technologies are advancing rapidly, the ability to engineer designer variations quickly and efficiently is essential. By manipulating large synthetic constructs >100 kb, we can study the relationship between genomic architecture and functional elements, and their association with developmental regulation, human disease and evolution. CREEPY with its simplex and multiplex editing capabilities, will directly benefit these studies and accelerate such engineering.

## Methods

### Yeast strains and culture

BY4741 was used to test chromosomal editing efficiency, with *ADE2* as the target. The yeast strain with *mSox2* construct (yLM1371) was also using BY4741 background. Yeast strains were grown using YPD as rich medium or defined SC with appropriate amino acids dropped out as selective media. All yeast transformations in this study were performed with standard LiAc/SS/PEG method^41^.

### CREEPY plasmids and gRNA assembly

The original Cas9 and gRNA expression modules were modified from p414-TEF1p-Cas9-CYC1t (Addgene# 43802) and p426-SNR52p-gRNA.CAN1.Y-SUP4t (Addgene# 43803). The *LEU2* marker was replaced with the *SpHIS5* marker by homologous recombination in yeast, generating the plasmid pCTC019. The Cas9 expression module (*TEF1* promoter, Cas9 CDS, *CYC1* terminator) and gRNA expression module (*SNR52* promoter, *Not*1 cutting site, *SUP61* terminator) were constructed using Gibson assembly into the pRS416 and pRS426 vectors, generating the single-plasmid system pYZ462 (Cas9/gRNA-CEN) and pYZ463 (Cas9/gRNA-2μ), respectively.

For the multiplex editing, a tRNA-gRNA array was used to express multiple gRNAs^23^. From pYZ463 (Cas9/gRNA-2μ), we first introduced one synonymous mutation (G to A, +171 from ATG) in Cas9 CDS to eliminate the *Bsm*BI recognition site with MISO^42^. Then, the tRNA module with a bacteria GFP was assembled to replace the original gRNA expression module (*SNR52* promoter)^43^. The final entry vector was pYZ960. The gRNAs were assembled as described before^23^. Briefly, the primers containing the corresponding gRNA sequences, adapters and *Bsm*BI recognition site were designed to amplify a DNA fragment with tRNA^Gly^ and gRNA scaffold. The PCR amplicons were gel purified and mixed with the entry vector. They were assembled together using Golden Gate assembly with *Bsm*BI^44^. The final plasmids were confirmed by Sanger sequencing. All the gRNAs used in this study were listed in Supplementary Table 3. The tRNA-gRNA array with gRNA.CTCF13, gRNA.CTCF17 and gRNA.CTCF25 was shown in Supplementary Table 7.

### Yeast transformations for genomic editing

In this study, 250 ng of Cas9/sgRNA plasmids were used in all the experiments to test the editing efficiency. The transformations were selected on plates with appropriate SC drop-out media, to select both CRISPR plasmids and *mSox2* construct. The plates were incubated for three days at 30ºC for single colonies, which were then re-streaked to another fresh plate with the same selective media to reduce background.

For all episomal editing experiments, 1 μg of donor template DNA for each target were transformed. The donor templates were PCR products with primers (Supplementary Table 4) amplified from corresponding gblocks (Supplementary Table 5). For all the targets, a 200 bp homologous arm was used at each end. The PCR products were purified with ZYMO DNA Clean and Concentrator column (ZYMO Research Cat# D4004). The concentration was measured by Qubit dsDNA HS kits with appropriate dilutions (Invitrogen Q32851), which was also double checked visually in an agarose gel where the density of target bands was compared with the DNA ladder (1 Kb Plus, NEB N0469S).

### Whole genome sequencing for episomal constructs

The yeast genomic DNA samples were prepared using a Norgen Biotek fungi/yeast genomic DNA isolation kit (Cat# 27300). The sequencing library was prepared using NEBNext Ultra II FS DNA library prep kit (NEB E7805L) with 500 ng genomic DNA as input. The whole genome sequencing was performed using an Illumina NextSeq 500 system using pair-end 36bp protocol. All raw reads were trimmed to remove adaptor sequence using Trimmomatic^45^, and subsequently mapped to original *mSox2* assembly and CRISPR construct as custom references, and mouse genome (mm10) using Bowtie2 software^46^, with Samtools^47^. The alignment was visualized using IGV (2.12.2) and Genome Browser.

## Data availability

All data and materials used in this study are available upon request. The CREEPY entry vectors (pYZ462, pYZ463 and pYZ960) have been deposited to Addgene (Supplementary Table 6). Their sequence files and maps are also available at Addgene.

## Acknowledgements

We thank members of the center for Synthetic Regulatory Genomics (SyRGe) at NYU Langone Health for general help and discussions. We thank Hannah Ashe, Gwen Ellis and Matthew Maurano for assistance with sequencing. We also thank Megan Hogan for helpful discussions about *mSox2* design and constructs. This work was supported by NIH/NHGRI grant 1RM1HG009491. The figures were created with BioRender.com.

## Author contributions

Y.Z. and J.D.B. conceived the project. Y.Z. and C.C. performed the experiments and collected data. Y.Z., C.C., S.L. and J.D.B. analyzed the results. J.M.L. built the pYTK plasmid. R.B. provided the wild-type *mSox2* assembly and designed most primers and gblocks used in this study. Y.Z., S.L., R.B. and J.D.B. prepared the manuscript with comments from other authors. This project was supervised by J.D.B.

