## Supplementary Figure for "Episomal editing of synthetic constructs in yeast using CRISPR"

### Slide 1
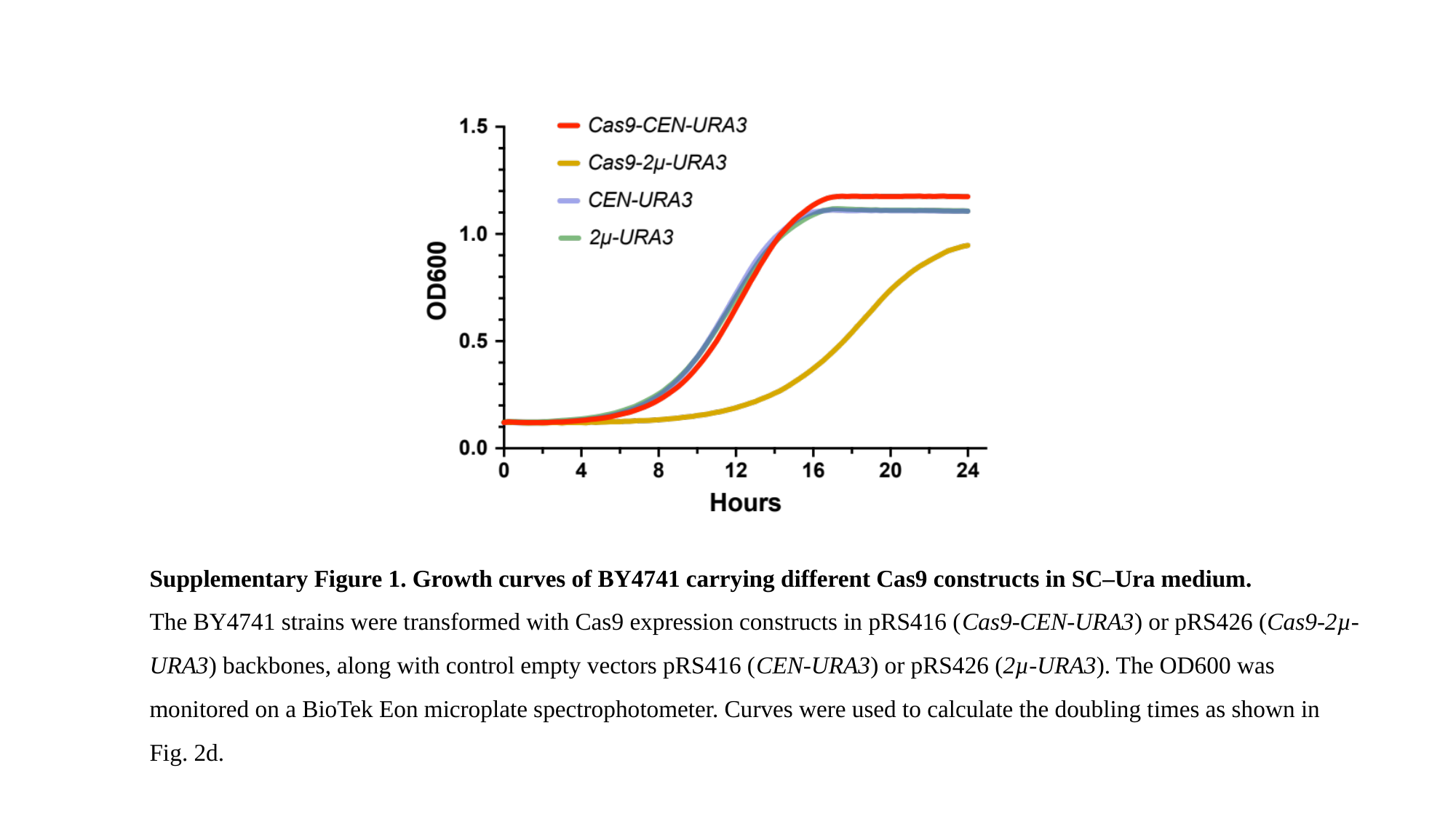

Supplementary Figure 1. Growth curves of BY4741 carrying different Cas9 constructs in SC–Ura medium.
The BY4741 strains were transformed with Cas9 expression constructs in pRS416 (Cas9-CEN-URA3) or pRS426 (Cas9-2µ-URA3) backbones, along with control empty vectors pRS416 (CEN-URA3) or pRS426 (2µ-URA3). The OD600 was monitored on a BioTek Eon microplate spectrophotometer. Curves were used to calculate the doubling times as shown in Fig. 2d.

### Slide 2
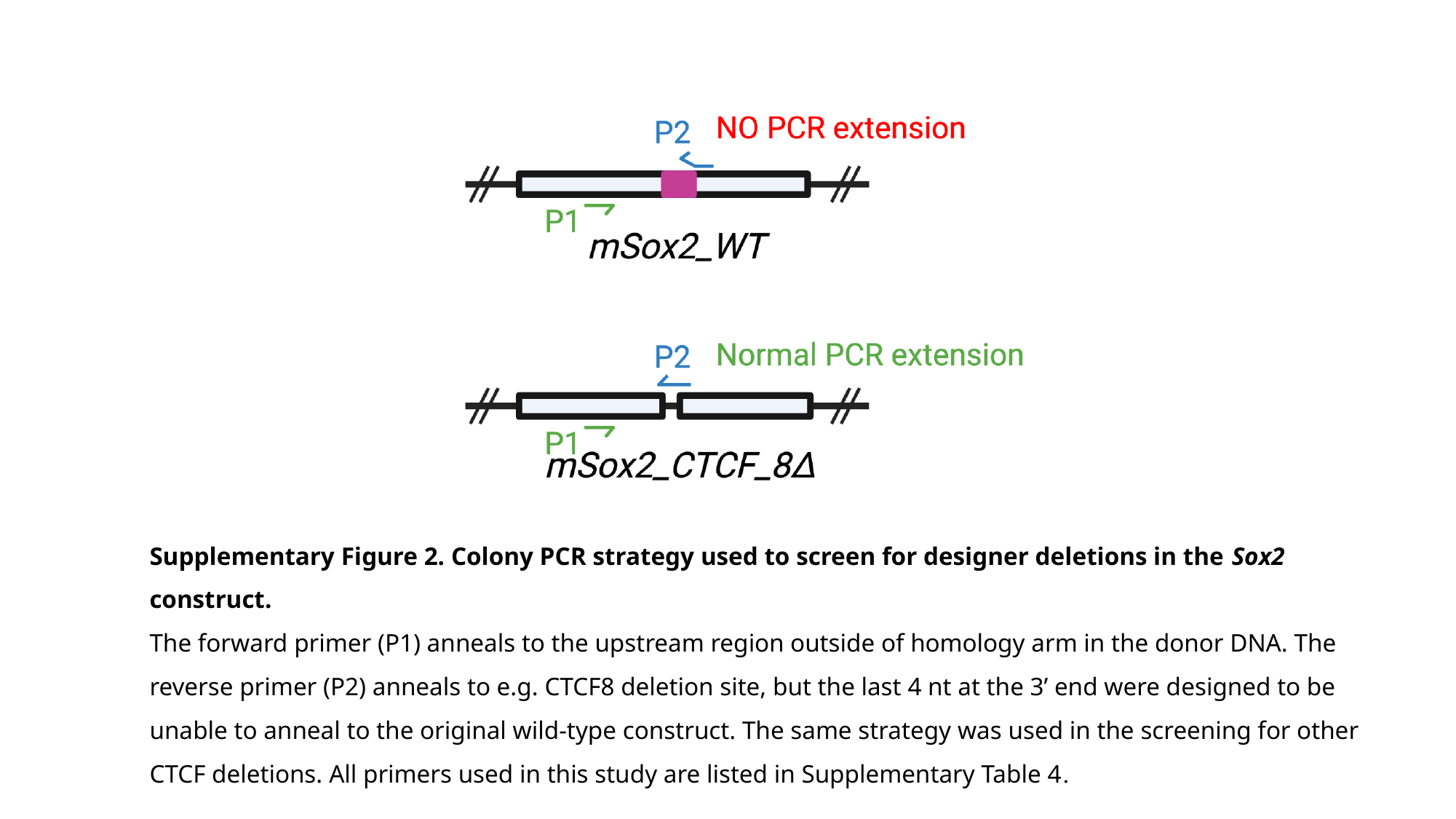

Supplementary Figure 2. Colony PCR strategy used to screen for designer deletions in the Sox2 construct.
The forward primer (P1) anneals to the upstream region outside of homology arm in the donor DNA. The reverse primer (P2) anneals to e.g. CTCF8 deletion site, but the last 4 nt at the 3’ end were designed to be unable to anneal to the original wild-type construct. The same strategy was used in the screening for other CTCF deletions. All primers used in this study are listed in Supplementary Table 4.

### Slide 3
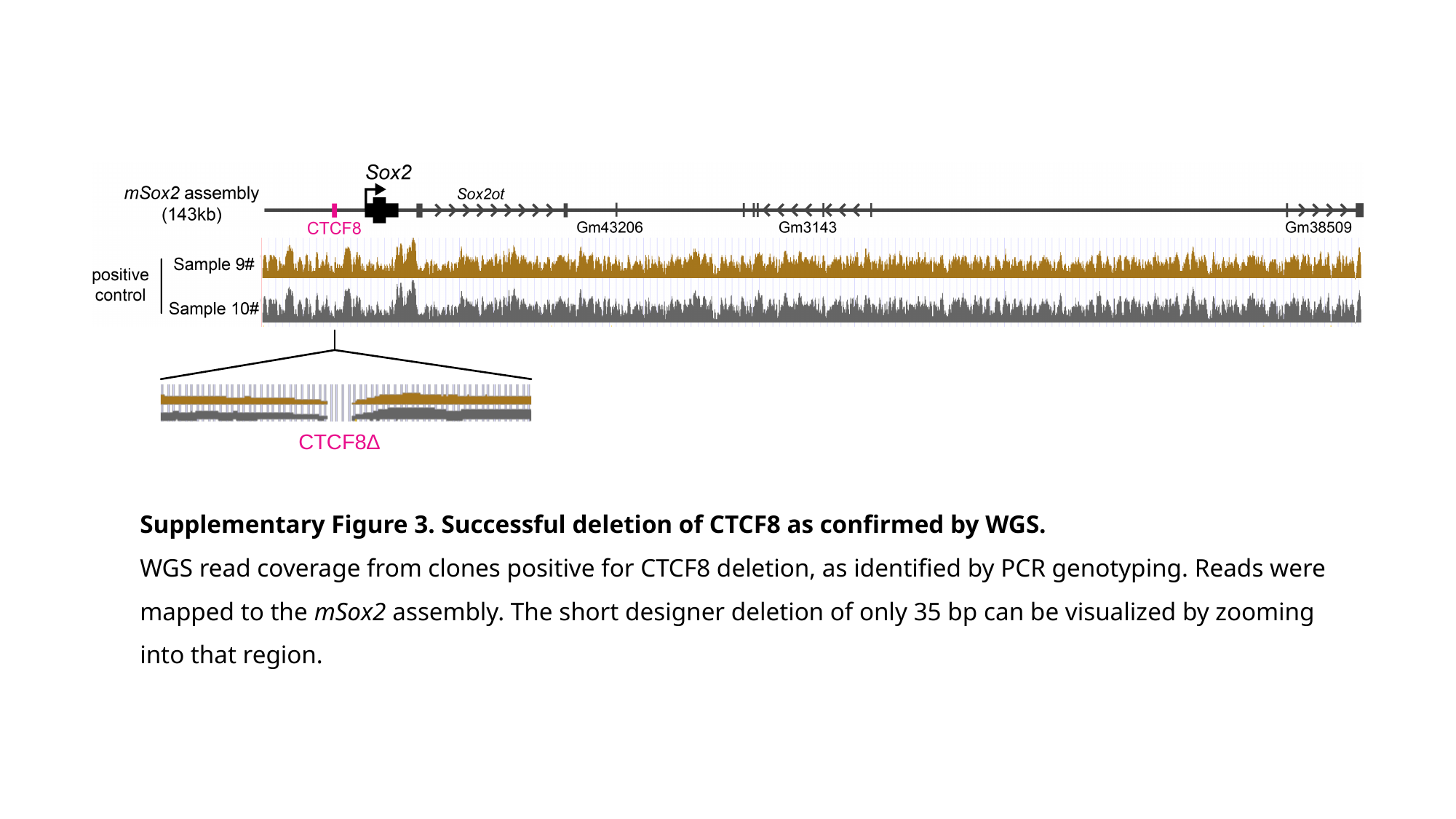

CTCF8∆
Supplementary Figure 3. Successful deletion of CTCF8 as confirmed by WGS.
WGS read coverage from clones positive for CTCF8 deletion, as identified by PCR genotyping. Reads were mapped to the mSox2 assembly. The short designer deletion of only 35 bp can be visualized by zooming into that region.

### Slide 4
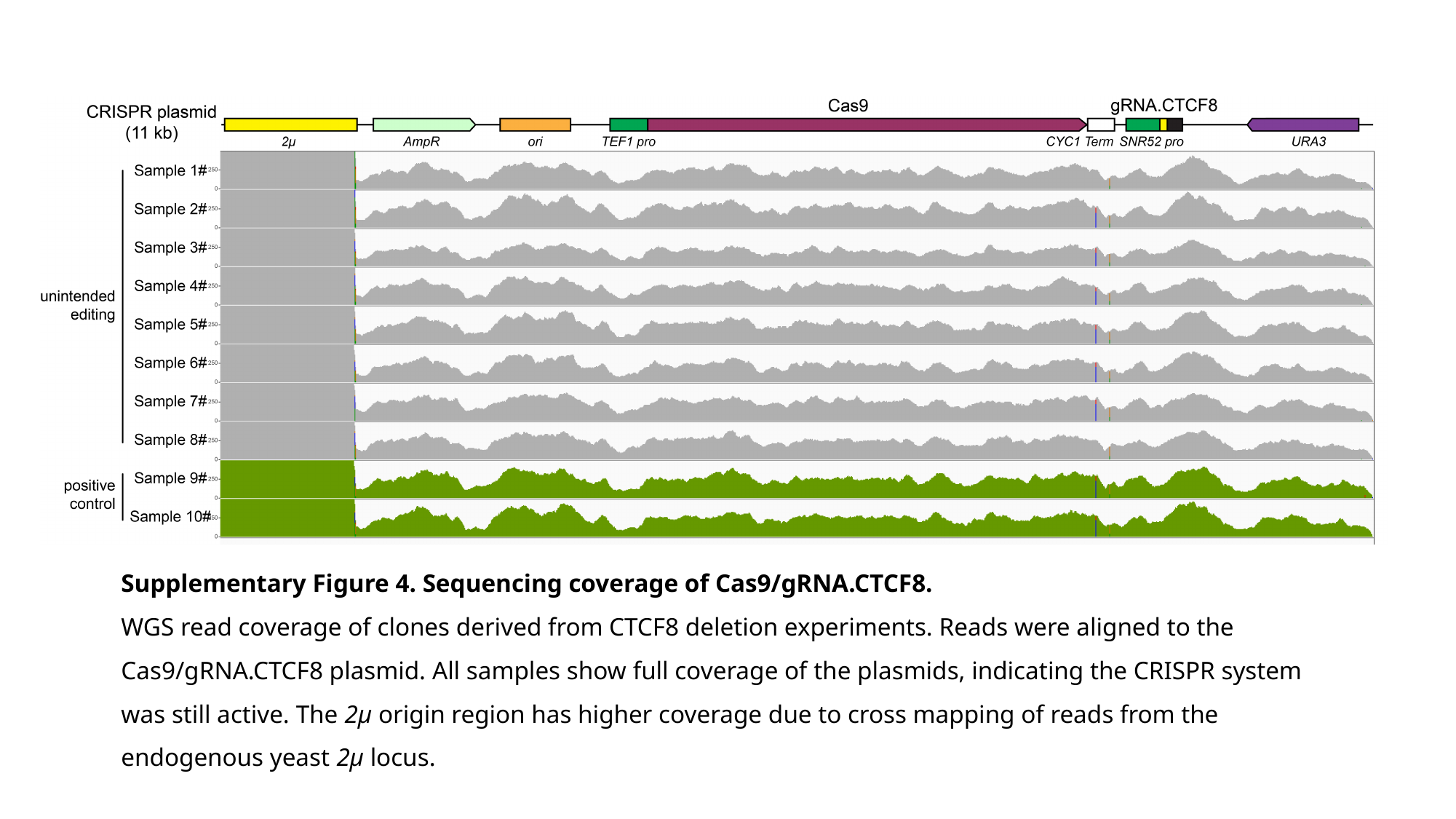

Supplementary Figure 4. Sequencing coverage of Cas9/gRNA.CTCF8.
WGS read coverage of clones derived from CTCF8 deletion experiments. Reads were aligned to the Cas9/gRNA.CTCF8 plasmid. All samples show full coverage of the plasmids, indicating the CRISPR system was still active. The 2µ origin region has higher coverage due to cross mapping of reads from the endogenous yeast 2µ locus.

### Slide 5
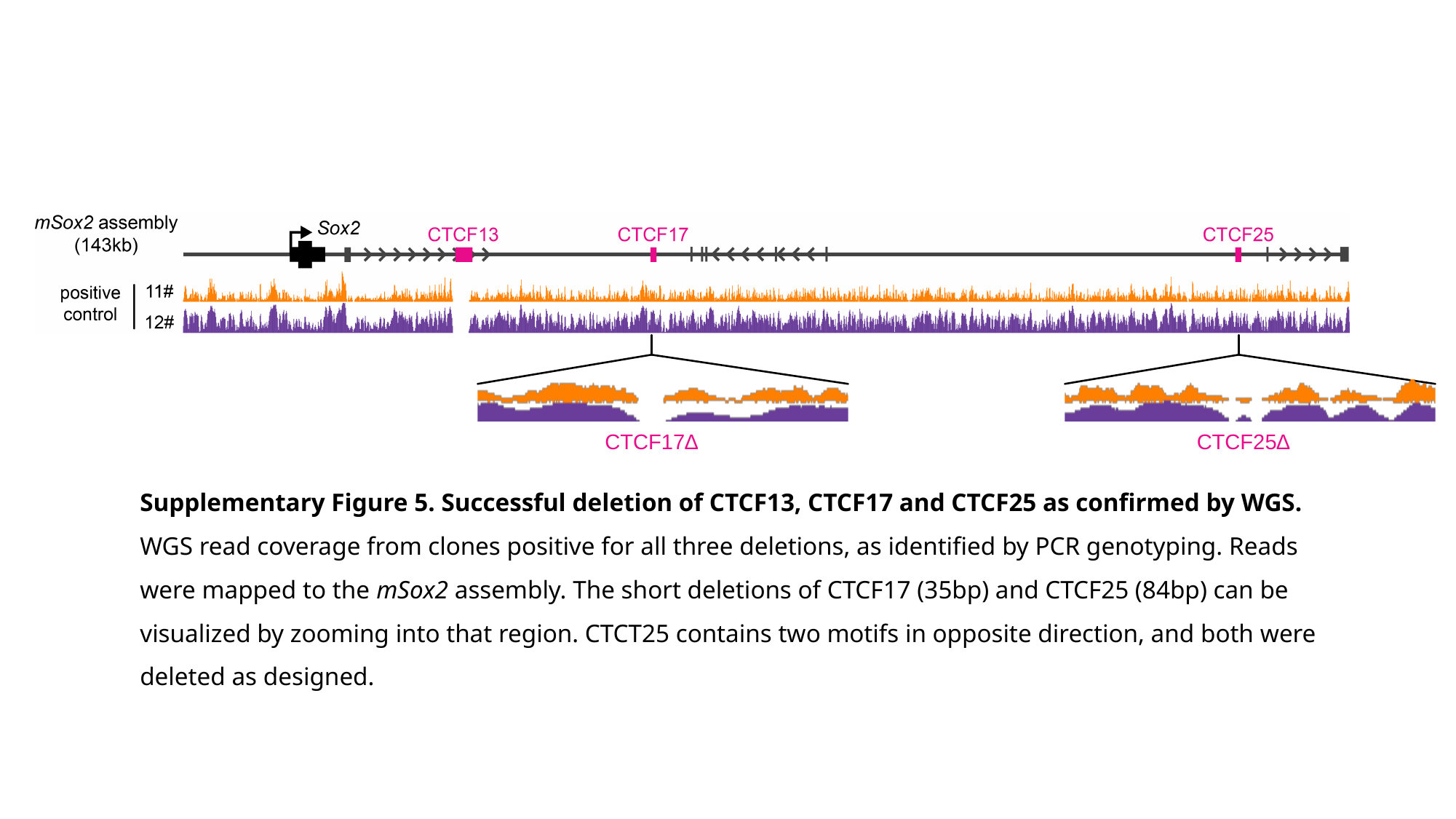

CTCF17∆
CTCF25∆
Supplementary Figure 5. Successful deletion of CTCF13, CTCF17 and CTCF25 as confirmed by WGS.
WGS read coverage from clones positive for all three deletions, as identified by PCR genotyping. Reads were mapped to the mSox2 assembly. The short deletions of CTCF17 (35bp) and CTCF25 (84bp) can be visualized by zooming into that region. CTCT25 contains two motifs in opposite direction, and both were deleted as designed.

### Slide 6
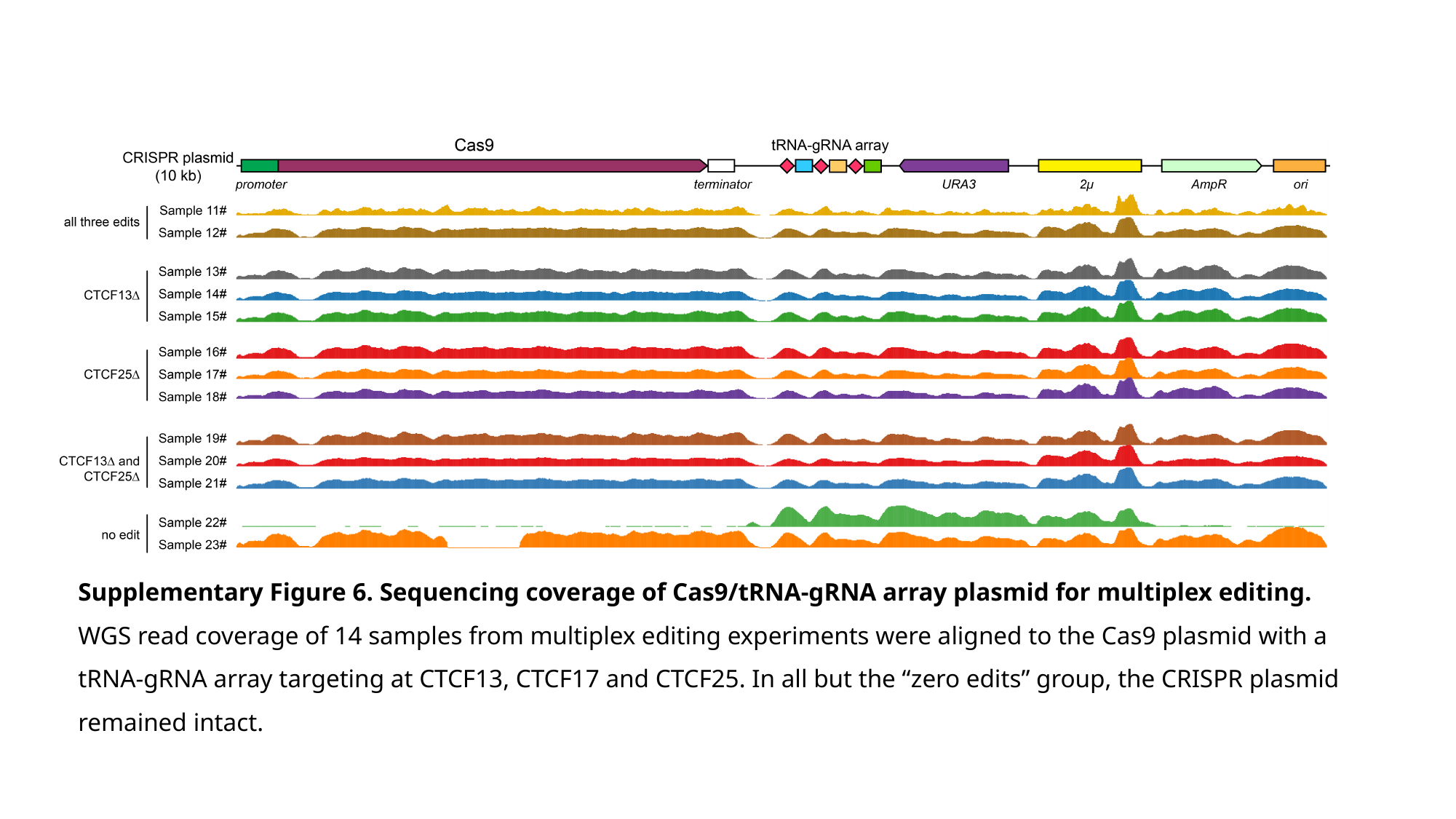

Supplementary Figure 6. Sequencing coverage of Cas9/tRNA-gRNA array plasmid for multiplex editing.
WGS read coverage of 14 samples from multiplex editing experiments were aligned to the Cas9 plasmid with a tRNA-gRNA array targeting at CTCF13, CTCF17 and CTCF25. In all but the “zero edits” group, the CRISPR plasmid remained intact.

### Slide 7
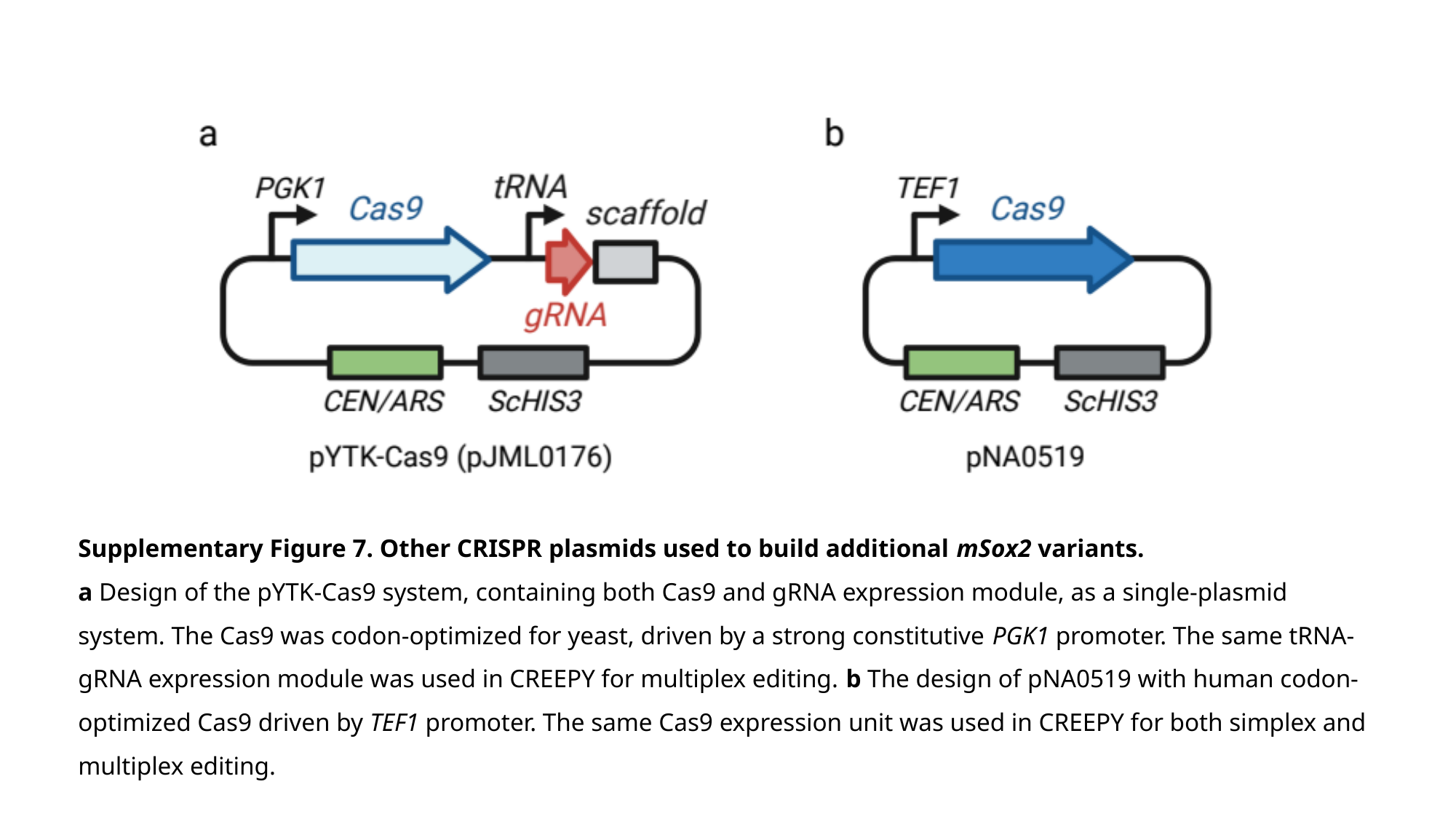

Supplementary Figure 7. Other CRISPR plasmids used to build additional mSox2 variants.
a Design of the pYTK-Cas9 system, containing both Cas9 and gRNA expression module, as a single-plasmid system. The Cas9 was codon-optimized for yeast, driven by a strong constitutive PGK1 promoter. The same tRNA-gRNA expression module was used in CREEPY for multiplex editing. b The design of pNA0519 with human codon-optimized Cas9 driven by TEF1 promoter. The same Cas9 expression unit was used in CREEPY for both simplex and multiplex editing.
