## Supplementary Table for "Episomal editing of synthetic constructs in yeast using CRISPR"

| sample | upstream deletion boundary | | | downstream deletion boundary | | | deletion length |
| --- | --- | --- | --- | --- | --- | --- | --- |
|  | repeat | type | coordinates (mm10) | repeat | type | coordinates (mm10) |  |
| 1 | micro-satellite | (TG)n | chr3:34638739-34638784 | micro-satellite | (CA)n | chr3:34667317-34667361 | 29 kb |
| 2 | micro-satellite | (GA)n | chr3:34636656-34636714 | Satellite | (GA)n | chr3:34668880-34669003 | 32 kb |
| 3 | SINE | B2_Mm1a | chr3:34633496-34633683 | SINE | B2_Mm1a | chr3:34706709-34706892 | 73 kb |
| 4 | micro-satellite | (GA)n | chr3:34636656-34636714 | Satellite | (GA)n | chr3:34668880-34669003 | 32 kb |
| 5 | SINE | B2_Mm1a | chr3:34633496-34633683 | SINE | B2_Mm2 | chr3:34707201-34707381 | 74 kb |
| 6 | SINE | B2_Mm1a | chr3:34633496-34633683 | SINE | B2_Mm1a | chr3:34706709-34706892 | 73 kb |
| 7 | micro-satellite | (CA)n | chr3:34635934-34635980 | SINE | B1_Mus2 | chr3:34690161-34690307 | 54 kb |
| 8 | low complexity repeats | CT-rich | chr3:34635398-34635702 | micro-satellite | (TTCC)n | chr3:34696689-34696761 | 61 kb |

**Supplementary Table 1. The repetitive elements to which deletion boundaries were mapped, from the simplex CTCF8 deletion experiment.**

The diagram representing these data is shown as Fig. 3b. These 8 samples are the same failed editing samples. The repetitive elements were found using RepeatMasker. The coordinates are the positions of the corresponding repeats as in mouse genome (mm10).

| edits | sample | upstream deletion boundary | | | downstream deletion boundary | | | deletion length |
| --- | --- | --- | --- | --- | --- | --- | --- | --- |
|  |  | repeat | type | coordinates (mm10) | repeat | type | coordinates (mm10) |  |
| CTCF13∆ | 14 | micro-satellites | (GAA)n | chr3:34680815-34680972 | micro-satellites | (GAA)n | chr3:34767804-34767978 | 87 kb |
| CTCF13∆ | 15 | micro-satellites | (TTCC)n | chr3:34684977-34685022 | micro-satellites | (TTCC)n | chr3:34761631-34761743 | 77 kb |
| CTCF13∆ | 16 | micro-satellites | (TTCC)n | chr3:34684977-34685022 | SINE | B1_Mus2 | chr3:34761896-34762031 | 77 kb |
| CTCF25∆ | 17 | micro-satellites | (TC)n | chr3:34638234-34638302 | SINE | B1_Mus2 | chr3:34689660-34689804 | 51 kb |
| CTCF25∆ | 18 | micro-satellites | (GA)n | chr3:34636656-34636714 | micro-satellites | (GAAA)n | chr3:34711313-34711491 | 75 kb |
| CTCF25∆ | 19 | micro-satellites | (GA)n | chr3:34636656-34636714 | micro-satellites | (TG)n | chr3:34737543-34737715 | 101 kb |
| CTCF13∆ and 25∆ | 20 | SINE | B1_Mus1 | chr3:34686574-34686723 | SINE | B1_Mus2 | chr3:34689660-34689804 | 3 kb |
| CTCF13∆ and 25∆ | 21 | SINE | B1_Mus1 | chr3:34686574-34686723 | SINE | B1_Mus2 | chr3:34689660-34689804 | 3 kb |
| CTCF13∆ and 25∆ | 22 | SINE | B1_Mus1 | chr3:34686574-34686723 | SINE | B1_Mus2 | chr3:34689660-34689804 | 3 kb |

**Supplementary Table 2. The repetitive elements to which deletion boundaries were mapped, from multiplex editing experiments.**

The diagram representing this data is shown as Fig. 4f. The same sample numbers are used here. The coordinates are the positions of the corresponding repeats as in mouse genome (mm10).

| gRNA sequence | PAM | target | source |
| --- | --- | --- | --- |
| GATATCAAGAGGATTGGAAA | AGG | *ADE2* | Zhang, et al (2019)^1^ |
| TAAAAGCAAGTCCACCAGCA | GGG | *Sox2_CTCF8* | this study |
| AGCTAGGAGCCCCTAGGTAC | TGG | *Sox2_CTCF13* | this study |
| GAACGCCACTTACTGCTTAC | TGG | *Sox2_CTCF17* | this study |
| CACAGAAGAAAAAGCGCGGC | AGG | *Sox2_CTCF25* | this study |

**Supplementary Table 3. All gRNA sequences and their targets used in this study.**

The gRNA.ADE2 targets at *ADE2* CDS and tested in previous study. The other gRNAs targeting at *Sox2* CTCF sites were selected based on available PAM sequence (NGG). All these gRNAs cut efficiently in yeast.

| primer name | primer sequence | anneals to | note |
| --- | --- | --- | --- |
| oMSH700 | TCCCACTTATCTGGCGGTTCTATG | gbMSH036 | PCR amplification for CTCF8∆ donor templates |
| oMSH701 | GTTGCTTTAGTTCTTGGGAACCCG | gbMSH036 | PCR amplification for CTCF8∆ donor templates |
| oMSH719 | GCGCGCGGACAATCATAATT | *mSox2* | Colony PCR primers to screen CTCF8∆ deletion |
| oRB427 | TTCTGAGATGTGGCTGGTGC | CTCF8∆ | Colony PCR primers to screen CTCF8∆ deletion |
| oMSH750 | GGCGCTGAGTGCAGAGAATTATC | gbMSH051 | PCR amplification for CTCF13∆ donor templates |
| oMSH751 | GCATCTTTTCCTTCTGGCTATGGTG | gbMSH051 | PCR amplification for CTCF13∆ donor templates |
| oMSH748 | GGGTTTAAAACGTGCACCCC | *mSox2* | Colony PCR primers to screen CTCF13∆ deletion |
| oMSH749 | GGGCAGTCTGAGCACAAGAA | CTCF13∆ | Colony PCR primers to screen CTCF13∆ deletion |
| oMSH702 | TCAGTGACTGATGTAGCAGGGG | gbMSH037 | PCR amplification for CTCF17∆ donor templates |
| oMSH703 | GTAGCCTTTCCCCCTAGATGCT | gbMSH037 | PCR amplification for CTCF17∆ donor templates |
| oRB426 | CCATCTGCAATGGGATCAGTG | *mSox2* | Colony PCR primers to screen CTCF17∆ deletion |
| oMSH729 | CCTCTCCAGGAAGTGTGCTG | CTCF17∆ | Colony PCR primers to screen CTCF17∆ deletion |
| oMSH731 | TCTTGCTGTGTAAGGCAGGG | gbRB005 | PCR amplification for CTCF25∆ donor templates |
| oMSH732 | ACAAAGGGAGAGTCTGGGGA | gbRB005 | PCR amplification for CTCF25∆ donor templates |
| oRB436 | CGCACAGAAGAAAAAGCGAGC | *mSox2* | Colony PCR primers to screen CTCF25∆ deletion |
| oRB437 | CCCCATTCGTTATATAAGTTTCTTGC | CTCF25∆ | Colony PCR primers to screen CTCF25∆ deletion |

**Supplementary Table 4. Primers used in this study.**

| name | usage | sequence |
| --- | --- | --- |
| gbMSH036 | CTCF8∆ | TCCCACTTATCTGGCGGTTCTATGTACATTTCTAAGTTAAGCAAATTAAATTTGATTCTTATGATTTCGTCCCACGGCTTATATGATATGTTAACTTTTCTTTTGTTTAGATTATGTGTAATTAGTTTGATCTTCCATCCTCCTCGGTTTTTTTTTTTTTAACTTTATTTTTACACACCTTCTTTTAAAGGTAAAAGCACCAGCCACATCTCAGAAACTAGGCGCGCTGAAAAGTCGGTGGCCGCCTTCCCTGGCTCAACCTTTGCTCTGGTCTCCAGAGATTCGTGTTGAGCGTAATAGTACACCCCGATTAAGGAGAAAACAATGCTTTAATTTGAAGAGTCCCAAATATTTACGCGTTTCTATAAGTCCTTTCCGGGTTCCCAAGAACTAAAGCAAC |
| gbMSH051 | CTCF13∆ | GGCGCTGAGTGCAGAGAATTATCTGGAGCGTGCTTGATCTGGGGCTGCGTAGTCAGCGTAGTCAGCTTCTGCTTTATTTGTTTGTTTGTAATCTCATATTTAGCAAGCCAGGACGGTGGCAGCGAGGGAGAAAGGAAGGGATGGAGGAGGGGGCTTCGCGCTAACCACACCCCTTCCTAATGAGTTAATTTCCATATTTGGGGGGCTCTTCACAGCAGCCCCCTTTGTTCTCCTGAGCTCCTCTCTGATTTCAAGCTTCACTCGCCTGCCCCCCCCCCAAGTTTGTGTGTTAATGACGCTATTACCATCTGGCTTGCCCTACTTTCCAAAAGATATAAATCTCCCAGCTTGACTGCCCACATTATCATTTAAGTGCCACCATAGCCAGAAGGAAAAGATGC |
| gbMSH037 | CTCF17∆ | TCAGTGACTGATGTAGCAGGGGCCAAGCCCACTGTGGCACTGTCAGGCTTAAGCAAGTAGATCCGGTTGCATAAACAGGCAAGTGAATGGGACGTAGCAATAGCTCATCCATCAGTTAAGAGCACCGCATGCTCTCCCAGAGGTCCTGAGTTCAATTCCCGGCAACCATATGGTGGCTCACAACCATCTGCAATGGGATCAGTGTACTTATATATGTAAAATAATTTTTTTTTAAAAAAGGGAGGAGCAAGTGAGTAAGCTGTGGAGAGCAAGGTAAGCAGTAAGTGGCGTTCCTCCACAGCCTCTGCCTCAGTTTCTACCCCAGGATTTTGCCCTCCACTCCTGCCTATTTCCTTTGATGATGGACTGTAAGCTGTAAGATGAAATAAACCCTTTCCTCACCAAATTGATTTTGATCATGGTGTTTTGTCATAGCAGAATAAATCTCATGTGTGTGTTTGTTATTAAGCATCTAGGGGGAAAGGCTAC |
| gbRB005*  (partial) | CTCF25∆ | TCTTGCTGTGTAAGGCAGGGTCTCAATAAGTGGTTCGTGCAGGCCCGGGGTTTCCTGATCTCTTGCCCAAGTGGCTGCTGTGCCCAGCATGTGGGCCCAATTTATTTTCCTACTTTTTCCAGGATGCGCAGATGCAGGAACTATCTTTGGTGTTTCTGTCTTTCTGCCTGCTCACCGCAGTTTCCGCTAACAAAACAAGCCTTTGTGCACTTTGTTTGCAGTGTTGAGAACTGATCACAGGGACGTGGGAGAACCCCTCCCTGGAGCCGCGCGCGGCGACCCTTTCTGTGGATCTGGCTTACTGAGGCTCAGTCCGCTTTGCCACACCGCCTCCTGCTGGCGGCTAGTTGACCTGCCGCGCTTTTTCTTCTGTGCGTCAGATACTTCACCAGGCCTGAGGGTCTTCTCTTTTCTTTTGAAATAGGGCTCAAGGAACCCAGGCAGACTGCAAACTCACTAAGTGGCCCGAAGCTGACTTTGAACTCATGATCTTCTTGCATCCATTTTCGTGGAGCTGGGGTTGCCCATGCATACCACCACACCCAGCTTTCCGAGCCAGATGAGGAGGGGGCTGCATTCTCAGACAACTCTCAGTGGCTAGCTTGCCCTTTGACAAAACTTTGCTAGGAATTTACCACCCCTACCCCCCCTCCCCAGACTCTCCCTTTGT |

**Supplementary Table 5. Donor templates used in this study to introduce CTCF deletions.**

These DNAs were first ordered as gblocks from IDT (Integrated DNA Technologies, Inc.). Then, they were amplified with PCR and purified for yeast transformation as donor templates. Red, upstream homology arm (~200bp); blue, downstream homology arm (~200bp). Underlined sequence, primer annealing sites for PCR amplification (Supplementary Table 4). * Only a part of gbRB005 (670 bp) is shown here, which contains the full-length donor templates used to delete CTCF25 in this study. The original gbRB005 is much longer (~1600bp), with more deletions for other CTCF sites that are not used in this study. The original gbRB005 was used in building extra *mSox2* variants^2^.

| plasmid name | note | yeast marker | yeast ori | antibiotics | Addgene # | Source |
| --- | --- | --- | --- | --- | --- | --- |
| p414 | p414-TEF1p-Cas9-CYC1t | *LEU2* | *CEN/ARS* | AmpR | 43802 | DiCarlo, et al (2013)^3^ |
| p426 | p426-SNR52p-gRNA.CAN1.Y-SUP4t | *URA3* | *2µ* | AmpR | 43803 | DiCarlo, et al (2013)^3^ |
| pYZ462 | TEF1p-Cas9-CYC1t and SNR52p-Not1-SUP4t | *URA3* | *CEN/ARS* | AmpR | 187970 | this study |
| pYZ463 | TEF1p-Cas9-CYC1t and SNR52p-Not1-SUP4t | *URA3* | *2µ* | AmpR | 187971 | this study |
| pYZ960 | TEF1p-Cas9(noBsmBI)-CYC1t and tRNA-GFP-SUP4t | *URA3* | *2µ* | AmpR | 187972 | this study |
| pYTK-Cas9 | PGK1-Cas9-PGK1t and tRNA-GFP-SUP4t | *ScHIS3* | *CEN/ARS* | KanR | N/A | this study |
| pNA0519 | TEF1p-Cas9-CYC1t | *ScHIS3* | *CEN/ARS* | AmpR | N/A | this study |

**Supplementary Table 6. Plasmids used in this study.**

Key entry vectors for CREEPY, pYZ462, pYZ463 and pYZ960 were deposited at Addgene. Their sequences and descriptions are also available at Addgene.

tRNA-gRNA array to express gRNA.CTCF13, gRNA.CTCF17 and gRNA.CTCF25

atgtgcttcagtattacattttttgccttcaacgccttgattgttctatttttgctaataataaatctatttcatcggactaaaagtccattagttgtaa**gcggatttagctcagttgggagagcgccagactgaagaaaaacttcggtcaagtcatctggaggtcctgtgttcgatccacagaattcgca**gatggccggcatggtcccagcctcctcgctggcgccggctgggcaacaccttcgggtggcgaatgggactttAGCTAGGAGCCCCTAGGTAC*gttttagagctagaaatagcaagttaaaataaggctagtccgttatcaacttgaaaaagtggcaccgagtcggtgc*aaacaagcgcaagtggtttagtggtaaaatccaacgttgccatcgttgggcccccggttcgattccgggcttgcgcaGAACGCCACTTACTGCTTAC*gttttagagctagaaatagcaagttaaaataaggctagtccgttatcaacttgaaaaagtggcaccgagtcggtgc*aaacaagcgcaagtggtttagtggtaaaatccaacgttgccatcgttgggcccccggttcgattccgggcttgcgcaCACAGAAGAAAAAGCGCGGC*gttttagagctagaaatagcaagttaaaataaggctagtccgttatcaacttgaaaaagtggcaccgagtcggtgc*tttttttattttttgtcactattgttatgtaaaatgccacctctgacagtatggaacgcaaacttctgtctagtggata

**Supplementary Table 7. The sequence of tRNA-gRNA array with gRNA.CTCF13, gRNA.CTCF17 and gRNA.CTCF25.**

**Lowercase bold underlined**, the first tRNA^Phe^ . Red lowercase, the tRNA^Gly^ genes in the array. Blue lowercase, HDV ribozyme sequence to promote editing efficiency at the 5’ end^4^. UPPERCASE underlined, the gRNAs for CTCF13, CTCF17 and CTCF25, from left to right. *Lowercase italic*, the gRNA scaffold downstream of gRNA sequence.

1.	Zhang Y*, et al.* A gRNA-tRNA array for CRISPR-Cas9 based rapid multiplexed genome editing in Saccharomyces cerevisiae. *Nature communications* **10**, 1-10 (2019).

2.	Brosh R*, et al.* Dissection of a complex enhancer cluster at the Sox2 locus. *bioRxiv* **2022.06.18.495832**, (2022).

3.	DiCarlo JE, Norville JE, Mali P, Rios X, Aach J, Church GM. Genome engineering in Saccharomyces cerevisiae using CRISPR-Cas systems. *Nucleic Acids Res* **41**, 4336-4343 (2013).

4.	Ryan OW*, et al.* Selection of chromosomal DNA libraries using a multiplex CRISPR system. *Elife* **3**, (2014).
